# *Fraxinus excelsior* updated long-read genome reveals the importance of MADS-box genes in tolerance mechanisms against ash dieback

**DOI:** 10.1101/2024.12.20.629733

**Authors:** Sara Franco Ortega, James A. Bedford, Sally R. James, Katherine Newling, Peter D. Ashton, David H. Boshier, Jo Clark, Susan E. Hartley, Andrea L. Harper

## Abstract

Ash dieback caused by the fungus *Hymenoscyphus fraxineus* has devastated the European ash tree population since it arrived in Europe in 1992. Great effort has been put into breeding programmes to increase the genetic diversity of ash trees and find heritable genetic markers associated with resistance, or tolerance mechanisms, to ash dieback. To facilitate identification of molecular markers, we used Oxford Nanopore Technologies combined with Illumina sequencing to obtain an accurate and contiguous ash genome. We used this genome to reanalyse transcriptome data from a Danish ash panel of 182 tree accessions. Using associative transcriptomics, we identified 175 gene expression markers (GEMs), including 11 genes annotated as dormancy MADS-box transcription factors which are associated with ash bud dormancy, flowering and senescence. We hypothesize that tolerant trees both break dormancy earlier in the year by increasing the expression of flowering-related SOC1 MADS-box and reducing the expression of SVP-like MADS-box, whilst also accelerating senescence by increasing the expression of JOINTLESS MADS-box genes. DNA methylation differences in the promoters of MADS-box genes between one tolerant and one susceptible tree indicate potential epigenetic regulation of these traits.

**Article Summary:** Ash dieback has devastated European ash tree populations. To aid in breeding programmes focused on finding solutions against this pathogen, we have assembled a new ash genome. This new genome helped us to identify genes related to tree biological life cycles, expressed differently in tolerant and susceptible trees. For the first time, we have also discovered that susceptible and tolerant trees showed different DNA methylation frequencies in those genes, suggesting epigenetic regulation. DNA methylation can turn on/off gene expression without changing the DNA sequence. These genes, and their regulatory elements, are ideal targets during breeding programmes combating this pathogen.

## Introduction

Common ash (*Fraxinus excelsior*) is a medium-sized deciduous tree with a wide distribution throughout all temperate zones of Europe. Ash is vulnerable to attack by many pests, for example, the emerald ash borer (*Agrilus planipennis*), a beetle native to Asia, whose larvae feed in the phloem, threatening the survival of the tree. However, since the first observation in 1992 in Poland, ash dieback caused by the ascomycete fungus *Hymenoscyphus fraxineus* (syn. *Chalara fraxinea*), has devastated European ash trees, killing over 90% of trees, reaching the UK in 2012 (Freer-Smith and Webber 2017). Ash dieback causes wilting and necrotic lesions on leaves, and later dot diamond-shaped lesions on the stems. Eventually, the disease kills the majority of trees, with only 5% of trees exhibiting low susceptibility to ash dieback. The teleomorph form of the fungi develops on the fallen rachises from leaves infected during the previous year and produces airborne ascospores reaching inocula of more than 100 spores m^3^, allowing transmission to other trees (Chandelier et al. 2014).

Due to both long generation and production cycles, forest tree breeding programmes emphasise the maintenance of genetic variability while selecting to improve particular traits (Neale and Kremer 2011). Genomics assisted breeding offers new opportunities to exploit wild tree populations whilst maintaining their genetic diversity (Migicovsky and Myles 2017). Genome-wide association studies (GWAS) can be used to determine loci responsible for a trait (resistance, tolerance or susceptibility) and their size effect on the phenotype. Previously, more than 3,000 single-nucleotide polymorphisms (SNPs), were associated with low vs. high ash dieback disease scores, and 61 were associated with the biotic stress response in other plant species. Prediction models using these data were able to estimate the health score of the tree with 90% accuracy (Stocks et al. 2019). However, this analysis was performed on a relatively low-contiguity ash genome reference (89,487 scaffolds with *N*_50_ of 104 kbp) (Sollars et al. 2016), which could cause an overestimation of marker-trait associations due to inaccuracies across regions with high numbers of undetermined nucleotides (∼17% of this genome reference).

Obtaining an accurate, contiguous genome is a key step for downstream analysis. Here, we report the assembly of a new high-contiguous genome for European ash, and an assessment of the genetic diversity in a Danish ash population with a high incidence of ash dieback (for more information about the panel, please refer to Harper et al., 2016). We used Oxford Nanopore Technologies (ONT) to generate long sequence reads, to obtain a more contiguous and complete ash genome. To assess the utility of this assembly, we analysed mRNA-Seq data via associative transcriptomics (AT) analysis, which identifies molecular markers associated with a trait measured across a diversity panel, and has been used previously to identify gene sequence and expression variants associated with ash dieback damage using a Danish population (n=182) (Harper et al. 2016). This analysis identified new genes associated with low and high disease tolerance to ash dieback and the updated gene annotations were subsequently used to perform phylogenetic analysis.

A great additional advantage of using ONT technologies is the possibility of calling methylated sites on the genome. In plants, cytosine DNA methylations (5mCs) occur in three sequence contexts CpG, CHG and CHH (where H=A, C or T). Methylation of cytosine plays an important role in biological processes, for example, in gene expression regulation, silencing of transposable elements, or stress response, as differential DNA methylation can be related to selective expression of defence-related genes (Arora et al. 2022). Understanding differences in methylation between tolerant and susceptible ash trees could help to understand the basis of these phenotypes. For the first time, we have linked the expression of genes associated with ash dieback tolerance to significant differences in the DNA methylation frequency of the promoters of gene expression markers identified by the AT analysis.

## Material and methods

### DNA extraction and sequencing

DNA was obtained from the leaves of a grafted *F. excelsior* tree which was used for an earlier ash genome assembly (isolate 2451S; Sollars et al., 2016), which was provided to us by Future Trees Trust. The parent tree located at Paradise Wood, Oxfordshire, UK, was produced by the self-pollination of a hermaphroditic *F. excelsior* tree growing in woodland in Gloucestershire, UK, (52.020592, −1.832804) in 2002 as part of the FRAXIGEN project (Fraxigen, 2005).

Leaves were ground to a powder in liquid nitrogen, before DNA was extracted using a high molecular weight gDNA extraction protocol. Briefly, ground leaf material is incubated with Calson buffer (100 mM Tris-HCl pH 9.5, 2% CTAB, 1.4M NaCl, 1% PEG 8000, 20 mM EDTA and 0.25% β-mercaptoethanol) containing proteinase K at 65°C for 30 minutes with intermittent mixing, followed by an addition of RNaseA for a further 30 minute incubation. An initial crude extraction was performed with 1 volume chloroform, and the DNA precipitated with 0.7 volumes isopropanol. The resultant DNA was then subject to a further purification using QIAGEN genomic-tips, according to the protocol developed by Vaillancourt and Buell, (n.d.). Extracted DNA was quantified with Qubit and Nanodrop prior to sequencing. Libraries were prepared using the SQK-LSK109 ligation sequencing kit, using extended incubations for nick repair, end preparation and ligation steps, and each sample run on a single PromethION R9.4.1 (FLO-PRO002) flow cell. Sequencing was performed on a PromethION 24 running MinKNOW software version 3.6.1, and highest accuracy basecalling performed using Guppy basecaller version Guppy basecalling software version 3.2.8 (ONT).

### Genome assembly

Initially, the chloroplast and mitochondrial genomes were assembled using CANU (version 2.1.1) (Koren et al. 2017) and polished using RACON version 1.4.20 and MEDAKA version 1.0.3. (Vaser et al. 2017) (Supplementary text 1). CANU was then used to assemble the genome with the remaining unmapped reads.

To curate the heterozygous genome, PURGE HAPLOTIGS (Roach et al. 2018) was used to reassign allelic contigs and obtain a haploid assembly. The curated assembly was plotted to calculate the coverage and find 0.5x and 1x coverage contigs using the histogram cut-offs: a low point of 3, a midpoint of 45 and a high cut-off of 390. All the contigs with abnormally high or low coverage according to the histogram were removed and those with an alignment score greater than 80% were also marked for reassignment as haplotigs (option -a 80 in the purge function). Previously published Illumina data (ERR1399574; Sollars et al., 2016) was used to polish the curated assembly using PILON version 1.24 (Walker et al. 2014).

Benchmarking Universal Single-Copy Orthologs (BUSCO version 4.0; Manni et al., 2021) was used to assess the completeness of the genome against the embryophyte and eudicot datasets. Statistical analyses were also performed with QUAST v 5.0.2 (Gurevich et al. 2013).

### Genome annotation

To annotate the genome, repetitive elements were identified using REPEATMODELER (version 2.1) and REPEATMASKER (version 2.1; Flynn et al., 2020). BRAKER (version 1.9; Brůna et al., 2021) was used to predict the gene models in the masked genome. Protein data from the viridiplantae (downloaded November 2021; https://v100.orthodb.org/download/odb10_plants_fasta.tar.gz; Zdobnov et al., 2021) was used in combination with the previously published mRNA-Seq data from roots (ERR1399494), cambium (ERR1399492), leaves (mother tree and 2461s, ERR1399495, ERR1399573) and flowers (ERR1399493). TSEBRA (Gabriel et al. 2021) with default configuration was used to combine the results of both proteins and mRNA-Seq predictions. Annotations were filtered by structure and function using GFACS (Version 1.1.2; Caballero and Wegrzyn, 2019) and ENTAP (Hart et al. 2020). EGGNOG (Huerta-Cepas et al., 2019;Cantalapiedra et al., 2021) was used for the functional characterization of the gene models. Transposable elements (TE) and tandem repeats were identified using the Extensive de-novo TE Annotator (EDTA; (Ou et al. 2019).

### DNA methylation

NANOPOLISH (v 0.13.2; Simpson et al., 2017) was used to identify methylated CpG sites. DNA regions with a cytosine followed by guanine from 5′ to 3′ direction were identified as well as the frequency of methylation in the whole genome. The frequency was only calculated when the log_lik_ratio (log like methylated - log like unmethylated) had a positive value, supporting methylation.

### Pseudochromosome construction

A closely related species with a chromosome-level genome assembly can be used as a reference to construct pseudochromosomes from the contigs. We used the chromosome-level *Fraxinus pennsylvanica* genome (Huff et al. 2022) in NTJOIN (version 4.2.1; Coombe et al., 2020) with k=32 and w=32, n=2 options. NUCMER 3.1 (Kurtz et al. 2004) was then used for mapping (options -c 500 -b 500 -l 100 -maxmatch) between the *F. pennsylvanica* chromosomes and the *F. excelsior* pseudochromosomes. Files were later filtered (delta-filter function) to only maintain the synteny regions with identity >=90% and with a minimum length of 100 bp.

Circlize package in R was used to create a chord graph of syntenic regions with identity >90% and using a minimum length of 100bp. A circular plot of the genome was then performed using the Circlize package in R, showing the position on the pseudochromosomes of the contig and genes, GC content, methylated frequency, coverage and LTR transposable elements.

Mapping between genes (using the coding sequence) of both species was carried out using the Synteny Imaging tool (SYNIMA; Farrer, 2017) using OrthoFinder.

### Population structure analysis of Danish trees

To investigate the genetic diversity of *F. excelsior* in Denmark, we used previously published data (n=182, Harper et al., 2016). The mRNA-Seq data was mapped using STAR 2.7.10 (Dobin et al. 2013) against the new genome assembly, and variant calling was performed using GATK version 4.2.4.0 using HaplotypeCaller. VCFTOOLS version 0.1.16 (Danecek et al. 2011) was used to merge variants for the individuals of each panel, and to filter out genotypes called below a minimum allele count = 5, minimum number of alleles = 2, maximum missing data 25% (across all individuals), SNPs with > 3 alleles and quality < 30. After filtering, the samples were merged using VCFTOOLS. Annotation of the variants was done with SnpEff version 5.1 (Cingolani, Platts, et al., 2012) and filtered with SnpSift (Cingolani, Patel, et al. 2012) to only keep SNPs located on exons. The vcf file was converted to geno format using VCFTOOLS and the ped2geno function (LEA package) in R. PSIKO v2 (Popescu and Huber 2015) was used to infer population stratification.

### Associative Transcriptomics (AT)

To assess the validity of the new genome, we repeated the AT analysis of ash dieback disease damage traits in the Danish panel (Harper et al., 2016). To identify any differences in the gene expression, TPMs (transcripts per million) of the mRNA-Seq data from the Danish panel were calculated using salmon (version 0.8; Patro et al., 2017). Salmon counts were input into R 4.3.3 with tximport (Soneson et al. 2016) to obtain TPMs. Regression analysis was then conducted to compare the expression (TPMs) of the 182 trees of the Danish panel against the ash dieback disease damage scores (for more information about how damage score was measured refer to Harper et al., 2016). The analysis included multiple test corrections using false discovery rate (FDR) with Benjamini-Hochberg adjustment and Bonferroni threshold to reduce the likelihood of identifying false positives. We identified gene-expression markers (GEMs) by setting FDR <= 0.05.

### Phylogeny of MADS-box genes

To identify the role of MADS-box genes, classified as GEMs, we retrieved the amino acid sequence of the MADS-box genes with FDR<0.05 in the AT analysis, and the sequence of another 63 MADS-box sequences from *Arabidopsis thaliana*, *Pyrus pyrifolia* and *Prunus mumus* downloaded from GenBank (NCBI, National Center for Biotechnology Information). Sequences were then aligned using MAFFT (version v7.490) (Katoh and Standley 2013), trimmed with TRIMAl (version 1.4 rev15; Capella-Gutiérrez et al., 2009) and the maximum likelihood phylogenetic tree was created using IQ-TREE (version 2.2.0-beta; Nguyen et al., 2015). Maximum likelihood was performed on 118 amino acids with the Q.insect+G4 chosen automatically according to the Bayesian Information Criterion. The Interactive Tree of Life online tool (I-TOL version 6.5.8) was used to visualise the tree.

### Methylation profile of tolerant and susceptible trees

With the aim of identifying potential epigenetic regulations that might control the expression of GEMs, we extracted DNA was using the method of Workman et al., (Workman et al. 2018) from a pair of Danish trees previously described in Harper et al. (Harper et al. 2016); association panel tree Ash66, and prediction panel tree DNASH35, which had ash dieback disease damage scores of 0 and 75% respectively. ONT library preparation, sequencing, basecalling and methylation calling were performed as described above. NANOPOLISH was used to call methylation as described above. Methylation was assessed across different genomic locales (gene bodies, 1 kbp upstream and 1 kbp downstream regions) of all the genes in the genome. Plots were generated using ggplot2 in R representing the mean methylation frequency (20 intervals each representing 5% of the length of the gene body or the mean in each of the 20 intervals representing 50 bp of the upstream and downstream regions). To identify potential epigenetic regulation of GEMs, we assessed the methylation frequency in the upstream region (promoters) of three GEMs with lower FDR (Fe_g19663, Fe_g22999 and Fe_g40353). We first removed outliers with identify_outliers function in R and then compared methylation frequency between the susceptible and tolerant tree to ash dieback using the function t_test in R. To associate DNA methylation changes with gene expression, we also assessed and plotted using ggscatter and reg.line in R, the expression of these three GEMS across the Danish panel.

## Results and discussion

### A high-quality *Fraxinus excelsior* genome assembly

We generated > 49 billion bp in 16,511,805 reads (41.1% Q20, 10.5% Q30) with an average read length of 2990.7, equating to > 47x coverage of the previously reported 877Mbp ash genome (Sollars et al. 2016). After chloroplast and mitochondrial assembly (Supplementary text 1), the remaining reads (14,785,500; 40 billion bases; 45.53x coverage) were assembled into 4353 contigs and, after polishing with Illumina reads, a genome with a total length of 866,153,998 bp, N50=336,336 (minimum length 1121 bp and maximum length 2,511,904) and a 34.22% average GC content was obtained. Although this assembly is 11Mbp smaller than the previous assembly (Sollars et al. 2016), we hypothesize that it is due to far fewer undetermined “N” bases in the new assembly (0% compared to 17%), which can lead to overestimation of genome size. The BUSCO completeness score against the eudicots database was 81.7%, consisting of 68.5% single-copy and 13.2% duplicated BUSCOs. Against the embryophyte database, the completeness score was 85.1% consisting of 71.6% single-copy and 13.5% duplicated (Fragmented:6.4%, Missing:8.5%, n:1614).

The genome was annotated using a combination of mRNA-Seq data and protein gene model predictions. In total 43,392 gene models were obtained. Out of these, 15,245 were mono-exonic and 28,147 multi-exonic with a median of 4 introns each. The median gene size, coding sequence size and exon size was 1181, 549 and 114bp, respectively. Functional annotations were obtained for 39,864 of those gene models. The number of gene models increased by 12% compared with the previous assembly (38,852 protein-coding genes (Sollars et al. 2016). Transposable elements accounted for 50.15% of the genome (previously reported as 35.9% in (Sollars et al. 2016)), with long terminal repeats almost 30% of the genome, terminal inverted repeats 13.8% and 6.5% classified as non-TIF (Table 1). The level of cytosine C5 methylation was 71% (median), with a similar percentage methylation in all pseudochromosomes and a median GC content of 43.4% (Figure S1B).

**Table 1.** Transposable elements (long terminal repeat, considering the 4353 contigs and genome size of 866153998 bp.

| Type | Class | Count | Number<br>of bases<br>masked | %masked |
| --- | --- | --- | --- | --- |
| LTR | Copia | 13,9021 | 1.14E+08 | 13.18% |
|  | Gypsy | 59,818 | 5.66E+07 | 6.54% |
|  | unknown | 130,001 | 8.78E+07 | 10.14% |
| TIR | CACTA | 73,048 | 2.44E+07 | 2.82% |
|  | Mutator | 158,792 | 4.92E+07 | 5.68% |
|  | PIF_Harbinger | 72,787 | 1.38E+07 | 1.59% |
|  | Tc1_Mariner | 14,873 | 4.39E+06 | 0.51% |
|  | hAT | 66,539 | 2.75E+07 | 3.18% |
| nonTIR | helitron | 157,970 | 5.65E+07 | 6.52% |
| total | interspersed | 872849 | 4.34E+08 | 50.15% |

**Table 2.** Pseudochromosome sizes and number of contigs (4353) were obtained in the new *F. excelsior* genome assembly.

| Scaffold | Pseudochromoso | Number | Size | Size |
| --- | --- | --- | --- | --- |

|  | me | of<br>contigs/s<br>caffold | pseudochromosome<br>considering only<br>contigs (bp) | pseudochrom<br>osomes (bp) |
| --- | --- | --- | --- | --- |
| ntJoin11 | Fe_Chr01 | 278 | 57,049,569 | 86,871,061 |
| ntJoin3 | Fe_Chr02 | 424 | 49,314,643 | 74,385,309 |
| ntJoin7 | Fe_Chr03 | 173 | 37,946,013 | 61,580,219 |
| ntJoin17 | Fe_Chr04 | 166 | 41,423,499 | 63,362,762 |
| ntjoin15 | Fe_Chr05 | 215 | 46,945,204 | 70,449,272 |
| ntJoin20 | Fe_Chr06 | 166 | 38,821,743 | 57,332,361 |
| ntJoin19 | Fe_Chr07 | 138 | 30,142,704 | 47,976,435 |
| ntjoin1 | Fe_Chr08 | 153 | 35,917,803 | 55,280,346 |
| ntJoin4 | Fe_Chr09 | 165 | 40,278,964 | 60,937,593 |
| ntJoin2 | Fe_Chr10 | 175 | 36,769,960 | 56,950,872 |
| ntJoin5 | Fe_Chr11 | 135 | 30,206,494 | 48,899,062 |
| ntJoin12 | Fe_Chr12 | 18 | 43,071,336 | 64,260,172 |
| ntJoin6 | Fe_Chr13 | 176 | 39,691,603 | 58,994,051 |
| ntjoin8 | Fe_Chr14 | 161 | 36,996,715 | 52,400,938 |
| ntJoin16 | Fe_Chr15 | 186 | 35,541,244 | 51,705,577 |
| ntJoin18 | Fe_Chr16 | 126 | 27,257,267 | 42,313,996 |
| ntJoin0 | Fe_Chr17 | 205 | 46,292,443 | 65,972,964 |
| ntJoin9 | Fe_Chr18 | 133 | 30,323,337 | 45,688,329 |
| ntJoin22 | Fe_Chr19 | 140 | 30,209,446 | 48,559,296 |
| ntJoin21 | Fe_Chr20 | 95 | 26,387,313 | 39,753,493 |
| ntJoin13 | Fe_Chr21 | 149 | 33,344,042 | 47,622,639 |
| ntJoin10 | Fe_Chr22 | 121 | 26,241,128 | 41,519,084 |
| ntJoin14 | Fe_Chr23 | 153 | 34,940,498 | 50,364,800 |
| Not-<br>assigned |  | 507 | 11,041,030 |  |

Using the *Fraxinus pennsylvanica* assembly (Huff et al. 2022), we assessed the synteny of the new *F. excelsior* genome and assigned the contigs into 23 pseudochromosomes covering 98.8% of the genome size (Figure S1A), but adding undetermined nucleotides to the assembly (N’s per 100 kbp = 33,878.9). Although some small contigs did not assemble within the pseudochromosomes, they only accounted for 1.2% of the total genome size and were likely discarded by NTJOIN which can occur due to too many sequence differences caused by repetitive contigs covering the same region of the genome or local misassembles (Table 1). A total of 19,063 genes were identified as orthologues between *F. pennsylvanica* and *F. excelsior* respectively (out of a total of 35,470 and 43,392 genes in the *F. pennsylvanica* and *F. excelsior* genomes respectively) (Table S1, Figure S2).

Overall, with the use of long-read sequencing technologies, we improved the contiguity and completeness of the previous ash genome based on a short-read assembly (Sollars et al. 2016) and reported a high level of transposable elements (>50%), methylation levels along the genome, and a 12% increase in the number of gene models. We also reported a chloroplast assembly with a larger size (192 kbp) than previous chloroplast genomes in other *Fraxinus* species such as *F. pennsylvanica*, which had a reported chloroplast genome of 155Kb (Yi et al., 2019). Chloroplast inverted repeats are usually misassembled, or the boundaries of these regions are not always accurately reconstructed: this could be the case with the chloroplast of the ash genome. Many problems can be also faced in the complex mitochondrial genome due to the large sizes and multiple dynamic structures including linear, branched or circular (Bendich 1996), which could explain the slight oversize of the mitochondrial genome (592 kbp) compared with the 581 kbp reported previously (Sollars et al. 2016).

### Population genetic analysis to unravel ash dieback tolerance mechanisms

We first mapped Illumina reads from the Danish population to the new genome reference (the average mapping of the Danish was 84.5%; Table S2). The Danish panel population structure was also assessed using PSIKO (Table S3), based on the membership coefficient (>60%).

A total of 175 GEMs with FDR <= 0.05 (Figure 1, Table S4) were identified using associative transcriptomics. Out of those, only Fe_g19663 passed the Bonferroni threshold, followed by Fe_g22999 and Fe_g40353 (Bonferroni > 0.05 and FDR < 0.05), all of which were identified as MADS-box genes. The gene previously reported as a cDNA-SNP marker for ash dieback susceptibility (Harper et al. 2016), had a high similarity (99%) with the CDS of the Fe_g18379 (FDR= 2.41E-05).

**Figure 1.**
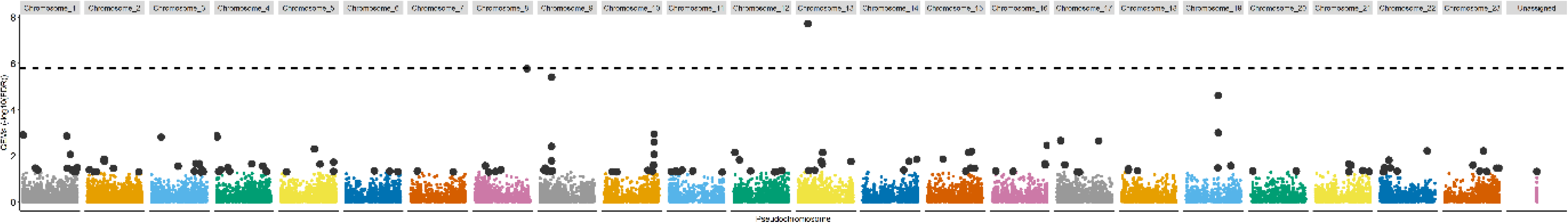
Gene expression markers (GEMs for ash dieback disease score) along the pseudochromosomes. The dotted line represents the Bonferroni cut-off, with only Fe_g19663 passing the threshold. Grey dots represent the GEMs with FDR<0.05.

A total of 11 MADS-box genes were identified within the 175 significantly associated GEMs (FDR<0.05). To better understand the role of these GEMs, we inferred a phylogenetic tree by creating a multi-locus sequence alignment with 63 sequences from *Arabidopsis thaliana*, *Pyrus pyrifolia* and *Prunus mumus,* obtaining a total of 118 amino acids after trimming. After the phylogenetic analysis, the GEMs were identified as likely orthologues of the SOC1-like MADS-box (Fe_g19663, Fe_g40350, and Fe_g40353, Fe_g40351.1), and the dormancy MADS-box JOINTLESS (Fe_g18379, Fe_g36254, Fe_g18374, Fe_g18375, Fe_g18373.1, Fe_g18376) (Figure 2). SOC1 (Suppressor of Overexpression of CO) has previously been identified as a promoter of flowering (Dorca-Fornell et al. 2011) whilst JOINTLESS has been associated with senescence (Nakano et al. 2015). Fe_g22999 was identified as S*HORT VEGETATIVE PHASE* (SVP)-like, a gene that in *Arabidopsis*, delays flowering by repressing floral regulators such as SOC1 (Li et al. 2008).

**Figure 2.**
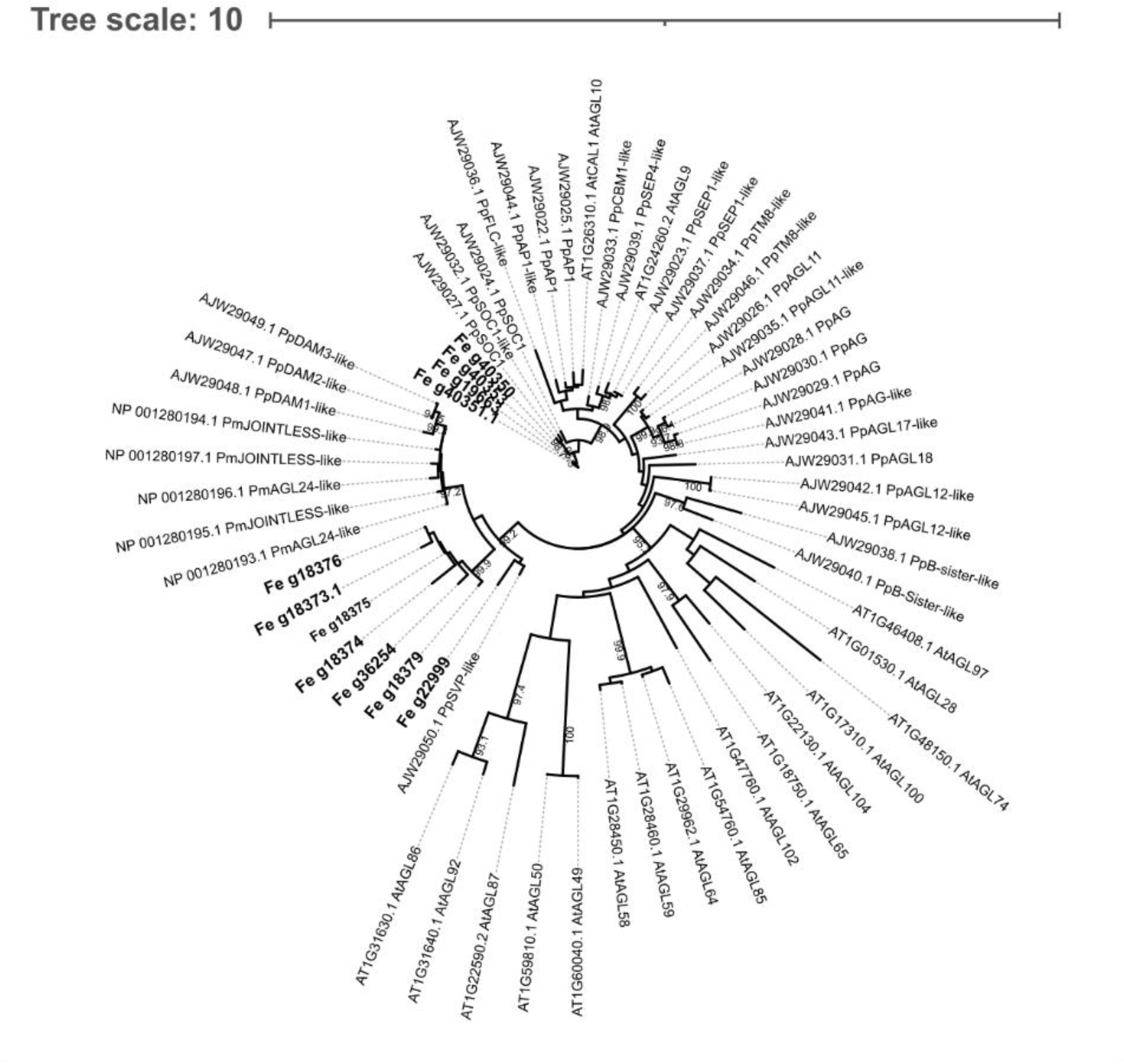
Maximum likelihood tree of the 11 MADS-box identified as GEMs. Bold genes are the MADS-box genes identified in the AT analysis and associated with susceptibility to ash dieback. The tree includes other MADS-box sequences from *Arabidopsis thaliana* (At)*, Pyrus pyrifolia* (Pp) *and Prunus mumus* (Pm) downloaded from GenBank (Accession number in the names). Bootstraps (from 90 to 100) are included in branches.

The high prevalence of MADS-box genes in the GEMs suggested a link between the phenology of the tree, such as dormancy, breaking-flowering and senescence, and susceptibility to ash dieback. In general, woody perennial plants modulate their seasonal phenology, growth and dormancy cycles, according to environmental conditions. Bud flush starts in the spring, and in the summer, the tree reaches the vegetative growth stage. In autumn, the trees stop growing and start forming buds, activating leaf senescence and abscission, whilst, in winter, endodormancy and cold acclimation occur (Ding and Nilsson 2016). After endodormancy, when the plant has accumulated enough chill, it gradually shifts to ecodormancy and if the conditions are favourable, the floral buds break (Alburquerque et al. 2008)(Wang et al. 2020). On the other hand, *Hymenoscyphus fraxineus* overwinters in the leaf litter on the ground and produces most of the spores from July to October, matching the vegetative growth stage of the ash trees.

Correlations between phenology and disease susceptibility have been already observed in other trees such *Quercus agrifolia* against *Phytophthora ramorum* (Dodd et al. 2008) or *Ulmus minor* trees against Dutch elm disease (Domínguez et al. 2022). Mckinney et al. (2014) found a moderate correlation in Danish and Sweden ash trees between earlier flushing and more resistant genotypes, which can explain this relationship with MADS-box genes. Therefore, we assessed if the different gene expression levels of the top three MADS-box (Fe_g19663, Fe_g22999 and Fe_g40353) correlated (Spearman correlation) with the damage score. We showed that the expression of SOC1-like Fe_g19663 and Fe_g40353, were negatively correlated with damage, whilst expression of Fe_g22999 was positively correlated (Figure 3). Considering the RNA samples were collected during the time of flushing (Harper et al. 2016), susceptible trees, with higher SVP-like expression might be expected to exhibit delayed phenology, whilst tolerant trees might flush and senesce earlier in the year due to reduced expression of SVP-like (Fe_g22999) and increasing expression of SOC1-like (Fe_g19663 and Fe_g40353) genes.

**Figure 3.**
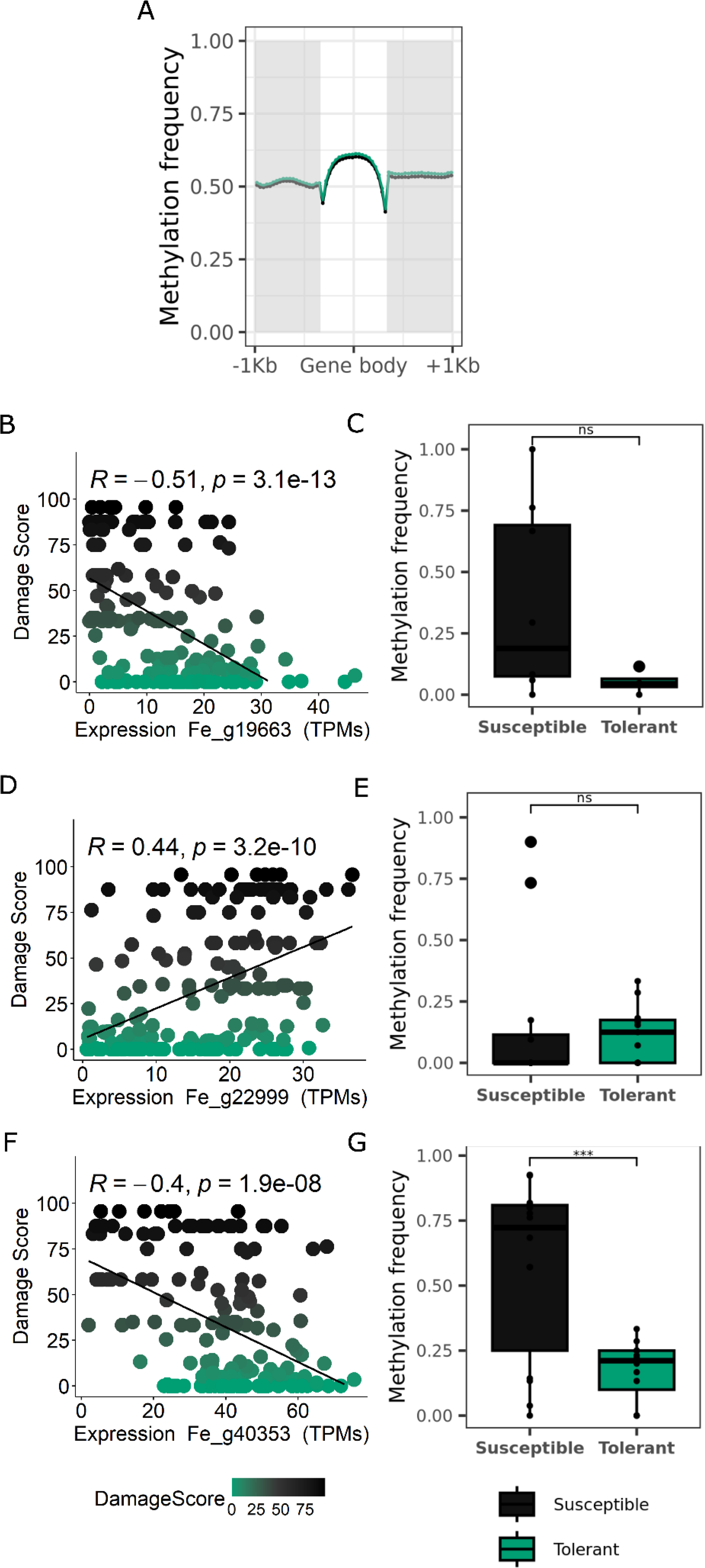
Expression and DNA methylation of genes in the new *F. excelsior* genome. **A**. Mean methylation frequency across all the genes in the new genome, according to the gene body and upstream and downstream regions (1 kbp each). Mean methylation was calculated in 20 intervals of the length of the gene body or the selected upstream and downstream regions. In green the results of tolerant tree to ash dieback and in black the susceptible tree. **B-D-E**. Spearman correlation between the expression of the GEMs Fe_g19663, Fe_g19663 and Fe_g40353, identified as SOC1, SVP-like and SOC1, respectively, and the damage score in each tree. The same colours as in A were applied. **C-E-G.** Methylation frequency differences between the susceptible and tolerant tree in the upstream region (1 kbp) of the GEMs: Fe_g19663 (p-value=0.054), Fe_g19663 (p-value=0.7) and Fe_g40353 (p-value=0.0008). Stars indicate significant differences. The same colours as in A were applied.

Epigenetic regulation of DORMANCY-ASSOCIATED MADS-BOX (DAMs) has also been reported, with DNA methylation found in the promoter of MADS associated with transcript repression (Rothkegel et al. 2017). However, until now, it has not been assessed whether there are differences in DNA methylation between susceptible and tolerant trees to ash dieback. As a preliminary investigation, we assessed DNA methylation in two trees with different degrees of susceptibility. More than 8.75 and 9.05 billion bases (35,0247,763 and 35,841,634 reads, N50=33797 and 25093 and Q20=14% and 15%) constituted the tolerant and susceptible ash samples, respectively. The coverage was >10x for each sample (Tolerant sample average read length was 248.3 and standard deviation=71,831.2; Susceptible sample average read length was 252.4 and standard deviation= 73,683.0). A total of 187,439,544 and 178,011,401 CpG sites and an average methylation frequency of 67.2% and 66.1% in the susceptible and tolerant trees, respectively. When looking at the methylation frequency across all genes (Figure 3A), no differences were seen between trees. However, we observed a significantly higher overall DNA methylation profile in the promoter of SOC1-like proteins in the susceptible tree (after removing outliers mean was 36.8% vs 5.2% in Fe_g19663 (p-value=0.0541 and 56.9% vs 17.3% in Fe_g40353 (p-value=0.00081) for susceptible and tolerant, respectively; Figure 3C, Figure 3G), which might result in lower expression of this gene (Figure 3B and Figure 3F). The opposite pattern was seen in SVP-like (Fe_g22999), with slightly but no significantly higher DNA methylation in the tolerant line (after removing outliers 15.8% vs 12.0% for susceptible and tolerant (p-value=0.7), respectively; Figure 3E), which again could result in lower expression of this gene (Figure 3D). Overall, the DNA methylation analyses showed variation in DNA methylation between a tolerant and susceptible tree, which consequently might regulate the expression of markers associated with different ash dieback susceptibility. However, to confirm epigenetic regulation of these genes, larger groups of susceptible and tolerant trees must be assessed.

## Conclusions

In summary, we have obtained a more contiguous ash genome which has allowed us to confirm the importance of phenology timing to combat ash dieback and identified 175 gene expression markers associated with disease damage scores. The new GEMs included several MADS-box genes, including SOC-1, SVP-like proteins and Dormancy MADS-box JOINTLESS, that showed differences in expression between susceptible and tolerant trees potentially regulated by differences in DNA methylation in the promoters. Identifying the loci responsible for tolerance to ash dieback could help in breeding programs of ash trees and combat this devastating pathogen.

## Data availability statement

The whole data of this project can be found under the BioProject PRJNA865134. Nanopore raw sequencing reads are available at NCBI SRA under the project number, SUB11822779 with the genome BioSample SAMN30100368, genome JANJPF000000000. Nanopore raw sequencing reads, and methylation profiles of the tolerant and susceptible samples are available in GSE214552, methylation profile of the new genome and salmon counts of the Danish population and are available at GEO under the number GSE214553 and GSE214554. Chloroplast and mitochondrial genomes are available at GenBank 2608965 and 2608833 and the sequence of the three MADS-box in GenBank with accessions (OP133138, OP133139 and OP133140). The new genome and annotations, the methylation profiles and the variant calling file can also be retrieved from https://webfiles.york.ac.uk/Harper/Fraxinus_excelsior/ and the scripts can be found at https://github.com/sfortega/Ash_genome.

## Acknowledgements

We gratefully thank Forest Research, Future Trees Trust and Earth Trust for allowing access to the land and plantations where the UK trees were sampled, and for providing tree grafts. We would like to acknowledge the Biological Technological Facility of the Department of Biology at the University of York for their technical support services along the project, especially to John W. Davey. We also want to thank the research computing cluster (Viking2) and the team supporting it, for allowing us to carry out all the bioinformatic analysis. We also gratefully thank the High-Throughput Genomics Group at the Wellcome Trust Centre for Human Genetics (funded by Wellcome Trust grant reference 090532/Z/09/Z) for the generation of sequencing data.

## Author contributions

SFO performed the genome assembly, annotation and methylation analysis, population mapping, associative transcriptomics analysis and wrote the first draft of this manuscript. JAB and ALH performed high molecular weight DNA extractions and generated a preliminary draft of the genome assembly. SJ prepared all the libraries, performed quality control and ran all the ONT runs; KN helped with the bioinformatic analysis and implemented the methylation analysis. PA supported the bioinformatic analysis, DB generated and JC maintained and grafted the reference tree, ALH, SHE and SFO planned the project and ALH coordinated project activities. SFO and ALH wrote the manuscript, and all authors reviewed and edited it.

## Supplementary

**Table S1.** Synima results using Orthofinder and the coding sequence of *F. excelsior* and *F. pennsylvanica* genes.

**Table S2.** Uniquely mapping rates using STAR of the reads of the Danish population.

**Table S3.** PSIKO population structure.

**Table S4.** Gene expression markers result with FDR adjusted.

---

**Figure S1.**
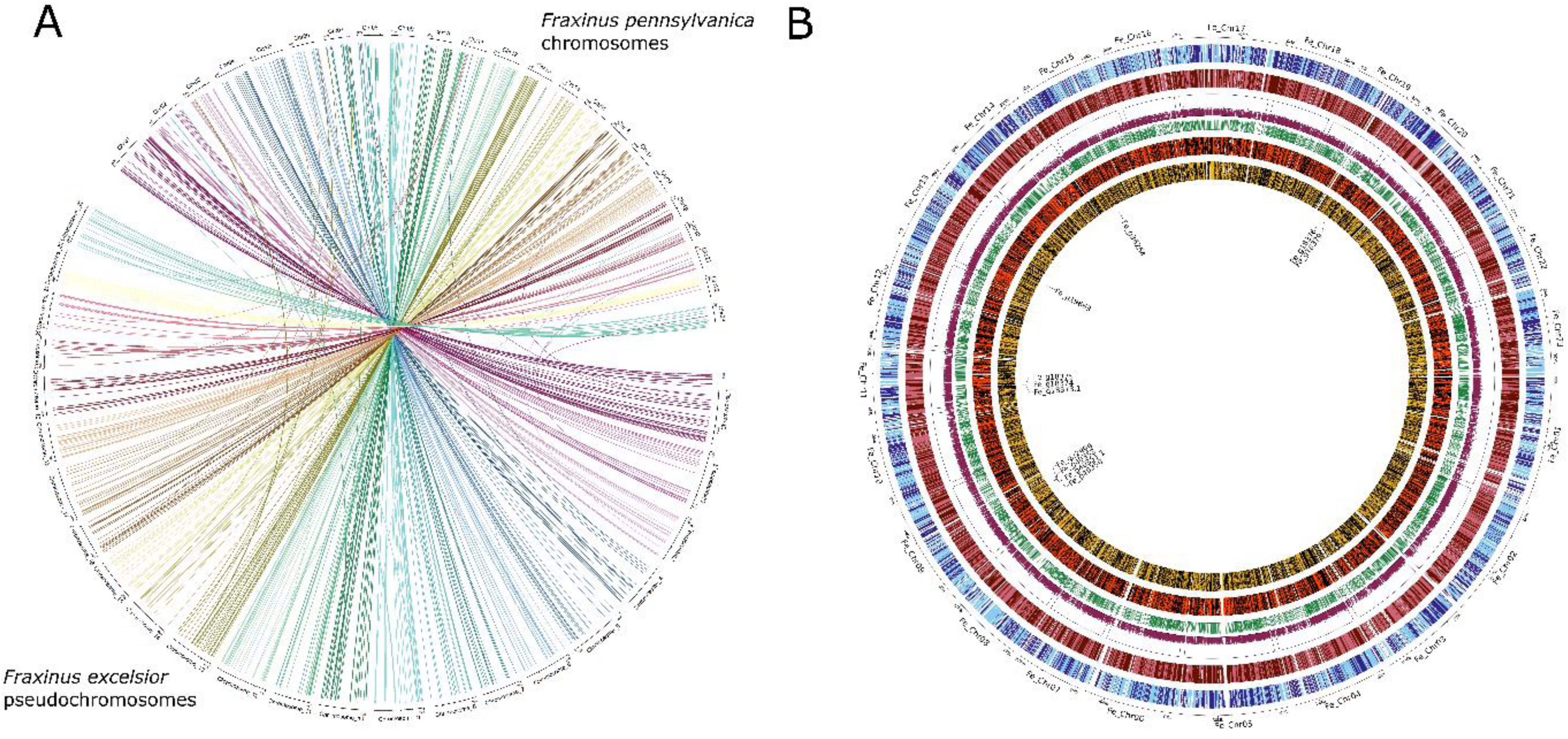
Plots of the new *F. excelsior* genome. **A**. Chord diagram showing similarity between *F. excelsior* contigs and *F. pennsylvanica* chromosomes. Each line represents regions with high similarity (>90%) between the sequence of *F. excelsior* contigs and *F. pennsylvanica* chromosomes. Each chromosome is represented in a different color. **B**. Plot of the F. excelsior genome. The circles represent (external to internal): contigs location (in the + strand in light blue and – strand in dark blue), genes (in the + strand in light red and – strand in dark red), GC content, methylation frequency, LTRs (in the + strand in orange and – strand in black), TIRs (in the + strand in yellow and – strand in black) and the MADS-box location.

**Figure S2.**
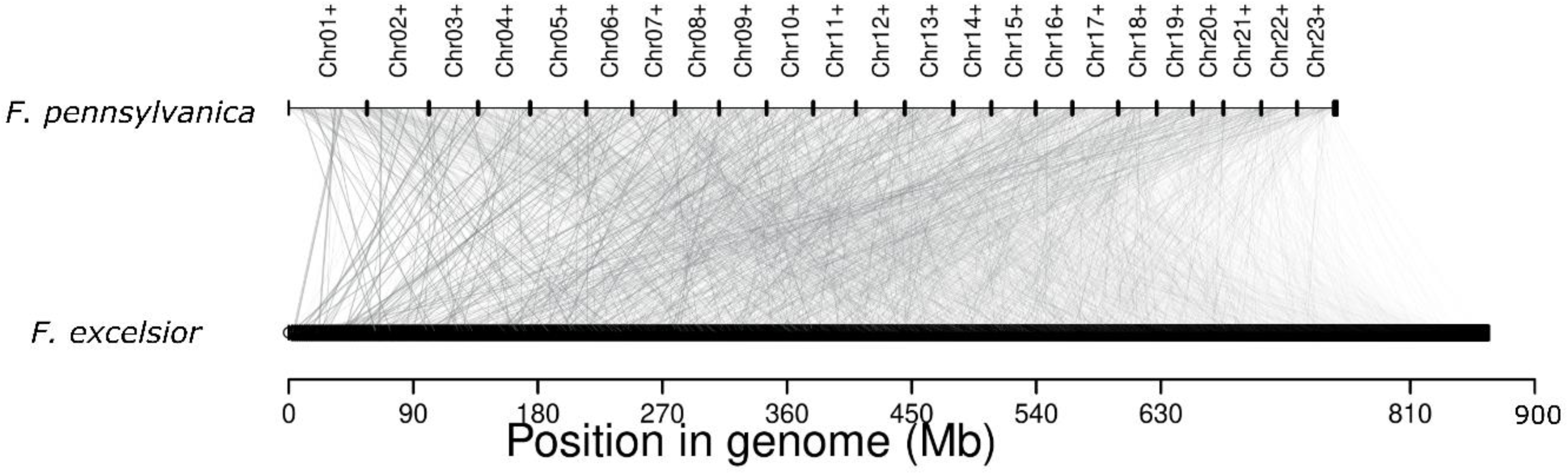
Gene synteny between *F. excelsior* and *F. pennsylvanica*.

